# Efficient laminar-distributed interactions and orientation selectivity in the mouse V1 cortical column

**DOI:** 10.1101/2024.11.05.621826

**Authors:** Licheng Zou, Giulia Moreni, Cyriel M. A. Pennartz, Jorge F. Mejias

## Abstract

The emergence of orientation selectivity in the visual cortex is a well-known phenomenon in neuroscience, but the details of such emergence and the role of different cortical layers and cell types, particularly in rodents which lack a topographical organization of orientation-selectivity (OS) properties, are less clear. To tackle this question, we use an existing biologically detailed model of the mouse V1 cortical column, which is constrained by existing connectivity data across cortical layers and between pyramidal, PV, SST and VIP cell types. Using this model as a basis, we implemented activity-dependent structural plasticity induced by stimulation with orientated drifting gratings, leading to a good match of tuning properties of pyramidal cells with experimentally observed OS laminar distribution, their evoked firing rate and tuning width. We then employed a mean-field model to uncover the role of co-tuned subnetworks in laminar signal propagation and explain the effects of intra- and inter-laminar coupling distributions. Our plasticity-induced modified model and mean-field model were able to explain both the excitatory enhancement through co-tuned subnetworks and inter-laminar disynaptic inhibition. Overall, our work highlights the importance of the clustering of neural selectivity features for effective excitatory transmission in cortical circuits.

## Introduction

Neurons in rodent primary visual cortex (V1) display spiking activity sharply tuned to the orientation of bars or edges in the visual field^1^. This property, termed orientation selectivity (OS), has been extensively studied in mammals^2–4^. Unlike cat and monkey, with V1 orientation-selective cells arranged in a semiregular, smoothly varying map across the cortical surface^2,5,6^, rodent V1 has no orientation map and neurons with different preferred orientations (PO) are intermixed^7,8^. Therefore, it remains an important focus in computational neuroscience to explain how the experimental results may be simulated accurately in terms of the orientation selectivity index (OSI), firing rates, tuning widths, without violating anatomical constraints^9^.

The anatomical pathway of visual signal transmission involves the lateral geniculate nucleus (LGN), which provides the principal thalamic inputs to V1 and was conventionally believed to convey only untuned inputs to the cortex. Consequently, OS was considered a visual feature computed in the cortex, beginning at the first stage of thalamocortical interactions^10^. Various computational models have been proposed to explain the appearance of OS in rodent V1, suggesting that the spatial arrangement of ON and OFF thalamocortical inputs to V1 accounts for the emergence of OS^11–14^. However, recent chronic two-photon calcium imaging studies suggest that axons of LGN neurons provide orientation- and direction-selective inputs to V1^15^ and this effect may be amplified by the recurrent circuit in V1^16^. This hints that at least some V1 neurons may inherit their preferred orientation (PO) from corresponding LGN presynaptic inputs and broadcast such selective information across the cortex. In addition, the recurrent nature of rodent primary visual cortex has been reported to follow a like-to-like organization principle^17–22^, indicating that neurons with the same PO have a higher probability to form reciprocal synapses. Such an organizational principle seems to be also relevant for various cognitive functions, such as working memory and decision making^23–27^. However, many computational models of PO have overlooked the potential contributions of cortical layers^28–34^ and cell-type-specific microcircuits^35^ to cortical computation. These elements are crucial for understanding inter-areal propagation^36– 38^, yet often omitted due to either a lack of experimental data or computational efficiency constraints. While the anatomical pathway of thalamic sensory input projecting to cortical layer 4 has been well characterized for the visual system, less is known as to how this information is processed and transformed within the cortex.

Given that the level of OS is significant across layers of the mouse V1 cortical column^1^, computational models can be employed to understand the extent to which the physiological findings described above account for the OS emergence^9^. On the basis of the latest anatomical studies on V1 inter-laminar organization and thalamocortical interactions, we built a biologically detailed model of the mouse V1 cortical column and implemented activity-dependent structural plasticity concerning new formation or deletion of synapses^39^ to induce corresponding like-to-like functional connectome where neurons with the same preferred orientation are more likely to interconnect with each other. The resulting tuning behavior of our network model in response to oriented drifting gratings matched existing experimental findings, including the laminar distribution of OS, evoked firing rate and tuning width of pyramidal cells. To decipher the role of co-tuned subnetworks in the process of laminar signal propagation, we utilized equivalent mean-field models to explain the modulation effects of cell-type specific stimulation on other neural populations determined by intra- and inter-laminar coupling distribution, and we found that this simplification could provide a mechanistic explanation of both the excitatory signal transmission enhancement through co-tuned subnetworks and the inter-laminar disynaptic inhibition upon stimulation on excitatory populations. Overall, our work suggests the importance of a feature clustering principle for effective excitatory transmission and provides a convenient way to analyze large-scale spiking networks.

## Results

We initially adopted the cortical column model (model #1) proposed by Moreni et al.^40^, which consisted of 5000 spiking neurons categorized into 17 different neuron types distributed across five laminar modules. Notably, layer 1 was exclusively constituted with VIP inhibitory neurons, while the remaining four laminae (layers 2/3, 4, 5 and 6) comprised pyramidal neurons as well as interneurons expressing parvalbumin (PV), somatostatin (SST) or vasointestinal peptide (VIP). Thus, a cell type was defined here by lamina as well as by neurochemical or expressed neuropeptide. Across layers with multiple neuron types, inhibitory populations were set considerably smaller than the respective excitatory populations, constituting 15% of the total number of neurons. The precise proportion of each inhibitory cell type in each layer was determined based on anatomical data^13^ and illustrated in Fig. 1A as the relative size of the respective inhibitory population^40^. The connectivity matrix adapted from previous work^13^ is shown in Suppl. Fig. S1.

**Figure 1:**
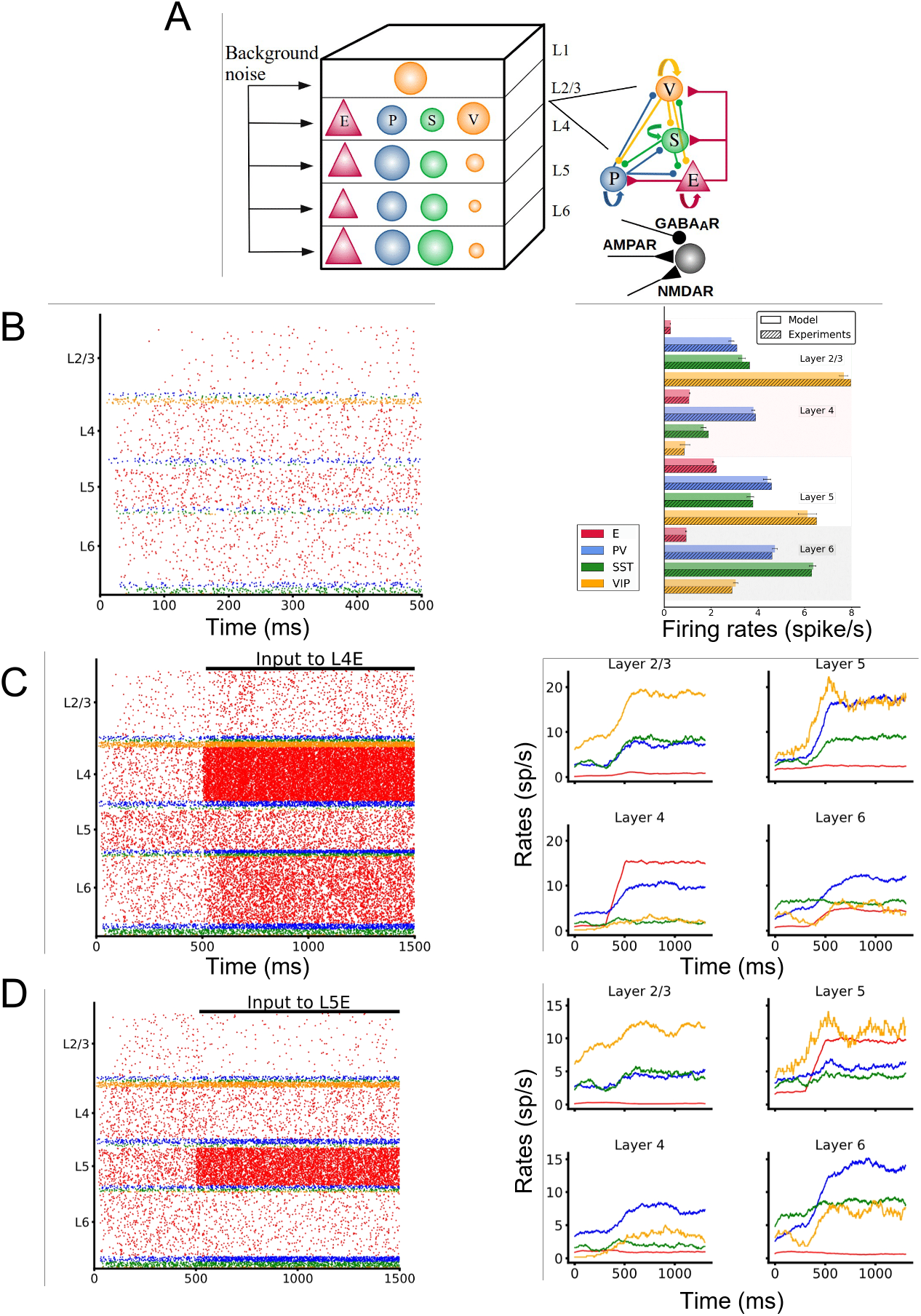
(A) Sketch of the cortical column model. In layers 2/3, 4, 5, and 6 an excitatory population E (red triangles) and three types of inhibitory population (PV, SST, VIP cells as blue, green, orange circles: P, S and V, respectively) are present. In layer 1 only VIP cells are present. The size of the circles in the top-left panel represents the relative size of the inhibitory populations. Connections between groups are not shown in the top left diagram; the zoomed-in schematic to the right shows inter-population connectivity and postsynaptic receptors (AMPA, GABA_A_, NMDA) involved. (B)-(D) Network in three different states. (B) Spontaneous (resting) condition. Simulated firing rates match experimental firing rates (right). (C) Feedforward (FF) condition (i.e., with excitatory input to L4 pyramidal cells). At 500ms a constant input of 30 pA is injected into all L4 pyramidal cells and excitatory neurons in all layers increase their activity. (D) Feedback (FB) condition (i.e., with excitatory 30 pA input into L5 pyramidal cells). At 500ms an input is injected in L5, and excitatory neurons in all other layers decrease their activities while interneurons in layer 6 significantly increase their firing.

### Three distinct states: spontaneous, feedforward-driven and feedback-driven

We started by examining three distinct states of the system: spontaneous (SP), feedforward-driven (FF), and feedback-driven (FB). In the SP condition, the cortical column received Poissonian background input across all cells (details provided in Moreni et al.^40^ Table S9), which were necessary to replicate the cell-type-specific firing rates (Fig. 1B) observed in vivo in mice^41,42^. When subjected to FF input, mainly deriving from LGN neurons projecting to layer 4 pyramidal neurons^15,16^, the system exhibited an increase in neuronal activity across all cell types throughout the various layers (Fig. 1C). However, when feedback signals representing top-down modulations from higher cortical areas were simulated by injecting currents into layer 5 pyramidal neurons, we found that feedback input concurrently diminished the firing rates of all other pyramidal neurons within the column (except for pyramidal neurons in layer 5 itself) (Fig. 1D). This phenomenon was attributed to the activity of inhibitory neurons, particularly in layers 4 and 6 (Fig. 1D right), which, in response to feedback input, substantially increased their firing rates, thereby inhibiting the pyramidal cells in non-layer 5 laminae. Consequently, the impact of feedback input contrasted with the effects observed with feedforward input.

### Activation of the cortical column through direction-selective input

We next introduced receptive field (RF) properties of neurons in the primary visual cortex by replacing constant FF currents with orientation-selective Poisson spiking neuronal inputs (Fig. 2A). In recent years, a plethora of literature has reported that the lateral geniculate nucleus (LGN) transmitted weakly selective inputs into V1^15,16,43–45^, and we implemented such input by modeling LGN neurons as Poisson neurons targeting layer 4 pyramidal neurons, with spiking frequencies following a Von-Mises distribution with direction as circular parameter (see Methods). Unlike the previous FF regime (Fig. 1C), selective inputs, presented under a fixed moving grating direction lasting for 1000 ms after a 500ms SP period, elicited responses from only a small population of layer 4 pyramidal neurons, with other neuron types showing less significant selective enhancement as their firing rates were subject to minimal changes (Fig. 2B). We then expanded the stimulus space across 16 different directions, evenly distributed from 0° to 360° (Suppl. Fig. S2A), with each selective input lasting for 500 ms. Within layer 4, visualization of the responses of a representative neuron unveiled both sharply tuned pyramidal neurons and more broadly tuned inhibitory neurons (Fig. 2C). Particularly for layer 4 pyramidal neurons, the sparse LGN connectivity probability (p=0.0015) resulted in extremely high OSIs for most neurons, with only a very few exceptions (Suppl. Fig. S2B).

**Figure 2:**
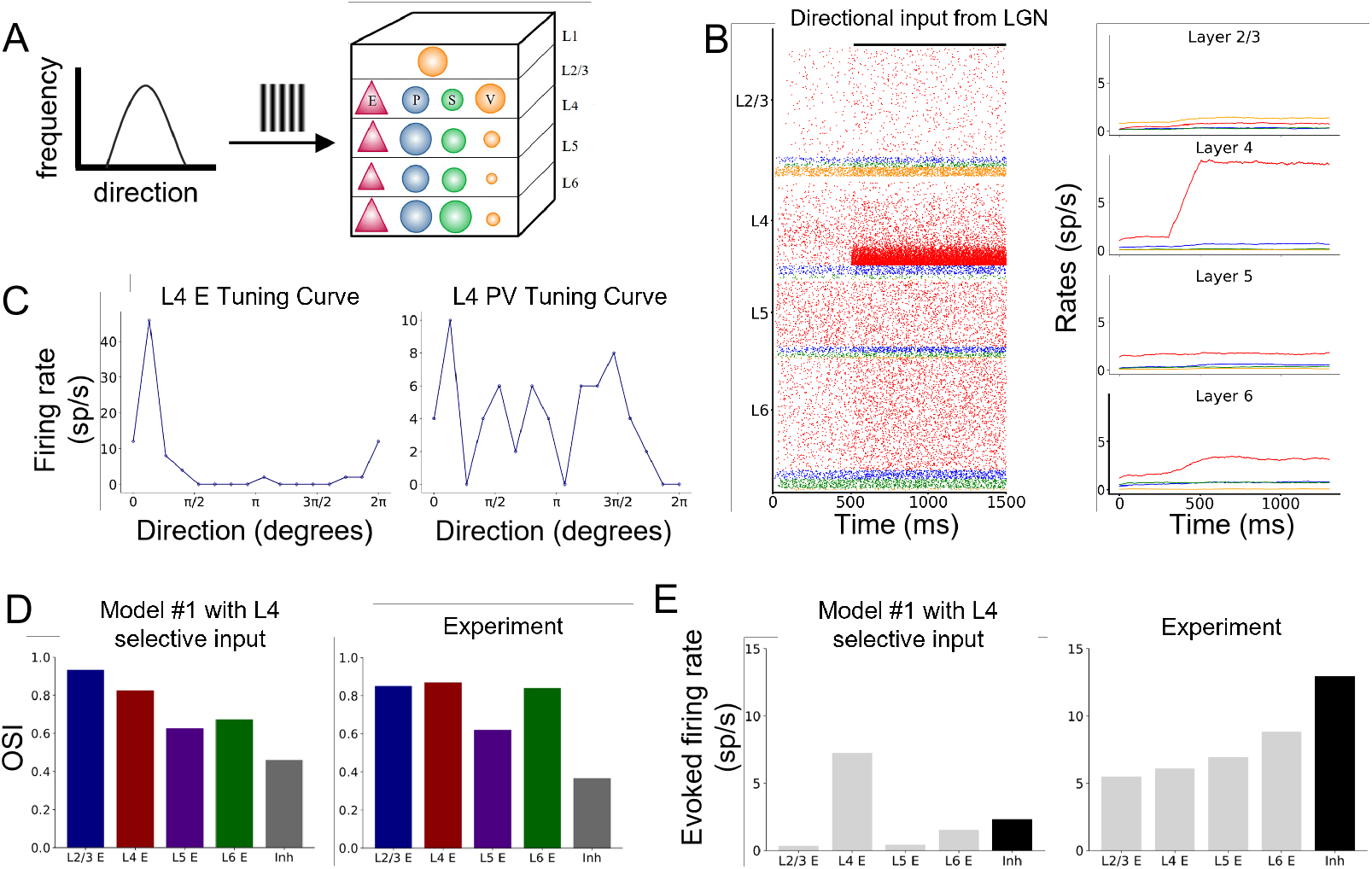
(A) Direction-selective project to layer 4 pyramidal neurons, illustrated for one visual direction. (B) At 500ms direction-selective inputs are sparsely injected to L4 pyramidal cells and representative excitatory neurons in all layers increase their activities. (C) After implementing direction-selective LGN inputs across 16 directions, the tuning curve of a L4 pyramidal neuron (left) and a L4 PV neuron (right). (D) Comparison of mean OSI in the model with experimental values^**1**^. (E) Comparison of mean evoked firing rate in the model with experimental values^**1**^.

We utilized the orientation selective index (OSI) as a metric to assess the level of OS (see Methods). We found that all excitatory populations in individual layers exhibited high OSI, with layer 5 displaying the lowest OSI and the other three layers showing high OSIs (Fig. 2D), consistent with a benchmark experimental study^1^. The high OSI in layer 4 pyramidal neurons stemmed from their reception of direction-selective LGN inputs. Signal transmission within the column, mainly from layer 4 to other layers, was not particularly strong, as indicated by our simulations (Fig. 2E). However, these limited presynaptic inputs from L4 to other layers, coupled with their low average firing rates, were sufficient to evoke orientation-selective postsynaptic responses (Suppl. Fig. S2C), leading to their higher than random (=0.5) OSI. For inhibitory neurons, the significantly higher baseline firing rates (Fig. 1C) together with densely connected synapses between pyramidal cells (Supply. Fig. S1) resulted in a higher denominator in the OSI calculation (see Methods), leading to low OSI scores.

Even though the OSI distribution closely mirrored data from the literature, the evoked firing rates varied significantly from the experimental data^1^, primarily because of the weak signal propagation behavior in our current model configuration (Fig. 2E).

### Recruitment of distributed thalamic FF and modulatory FB transmission for evoked firing rates

To fit both the OSI distribution and the evoked firing rates to existing experimental data^1^ at the same time, we designed a model configuration where both thalamic FF and cortical FB were considered. Specifically, thalamic FF inputs (which include LGN projections to layers 4 and 6 of V1 but also projections from other regions to all V1 layers) were modeled as distributed selective Poisson neuronal inputs targeting pyramidal neurons across all layers, while constant FB currents were injected on various interneuron types in deep layers (Fig. 3A). This network setup was based on evidence showing that under realistic conditions, rodent cortical columns in V1 often operate under a mixed regime, receiving both sensory feedforward inputs conveying external information and modulatory feedback signals originating from higher cortical areas simultaneously^26,27,38^. In this setup, thalamic FF axons would project to the entire cortical columns in V1^45^, while FB modulation from higher sensory areas would be distributed across deep layers but avoid layer 4^36,37^. It should be noted that the FF and FB configuration chosen here was designed to fit the experimental data rather than accurately capturing the realistic axonal projections found in literature.

**Figure 3:**
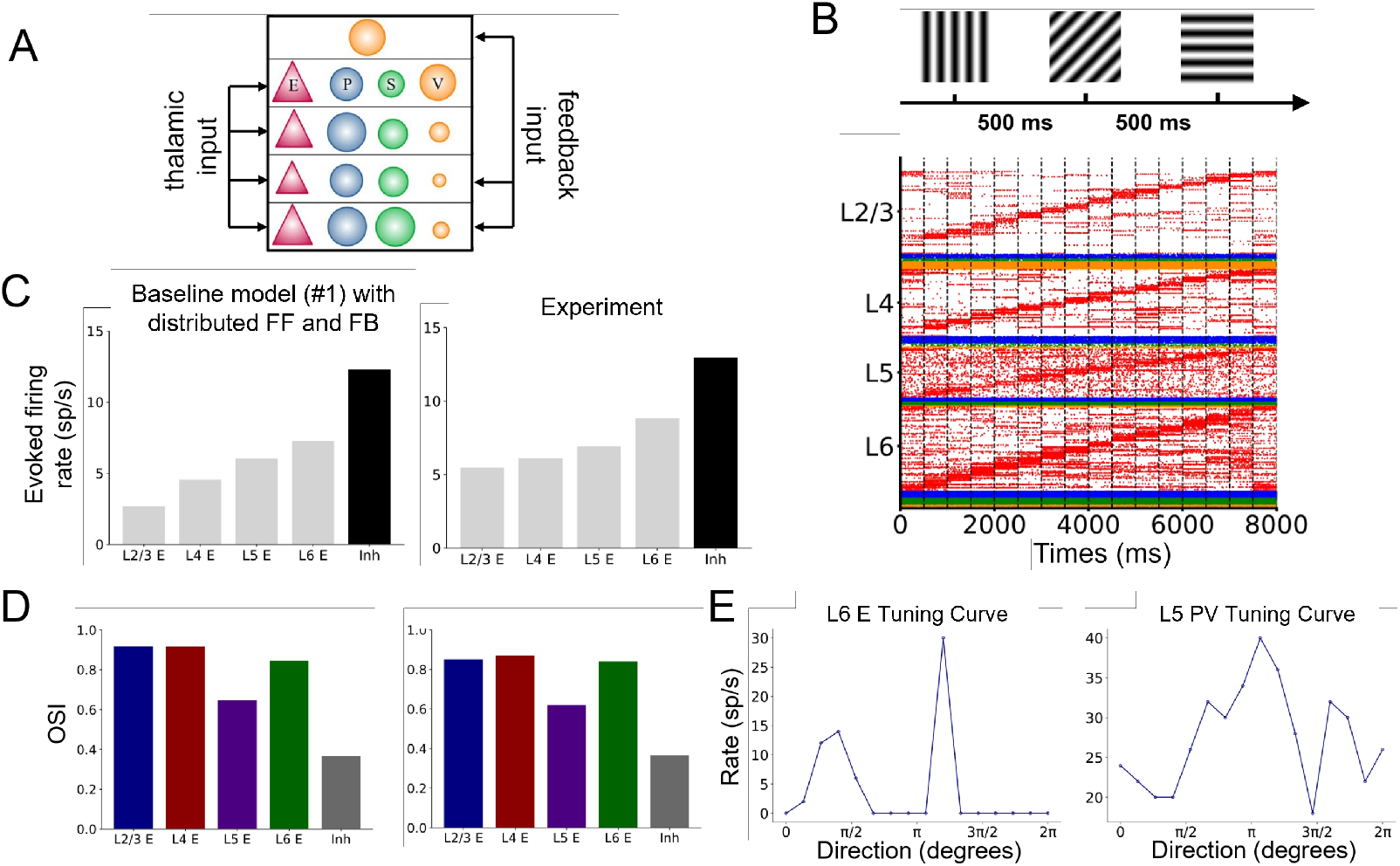
(A) Distributed LGN selective inputs project to pyramidal neurons across laminar layers and FB inputs are injected to layer 1, layer 5 and layer 6. (B) Each visual orientation lasts for 500 ms and firing rates of LGN Poisson neurons follow the Von Mises distribution. Raster plots of the column responses to orientational stimuli, reordered according to individual firing rates at each orientation bin. (C) Comparison of mean evoked firing rate in the model with experimental values^**1**^. (D) Comparison of mean OSI in the model with experimental values^**1**^. (E) After implementing selective inputs across 16 directions, the tuning curve of a L6 pyramidal neuron (left) and a L5 PV neuron (right).

Following the same simulation protocol as depicted in Suppl. Fig. S2, we exposed the cortical column model to 16 different visual directions, with each stimulus lasting for 500 ms (Fig. 3B). We found a highly selective behavior in pyramidal neurons when reordering their firing rate responses according to orientation bins (Fig. 3B). Comparing the mean evoked firing rate in the model with experimental values^1^, this time there was a good agreement between simulation results and experimental data, as all layers had received sufficient external input to drive the firing rates up (Fig. 3C). The same held true for the OSI distribution across all layers (Fig. 3D), with OSI values in excitatory populations remaining significantly high in layer 2/3 and layer 4, and increasing in layer 6 compared to the previous case (Fig. 2D). This OSI increase of layer 6 pyramidal cells was the direct consequence of them receiving direction-selective Poisson neuronal projections, resulting in higher OSI values even in the presence of strong evoked firing rate responses. However, layer 5 pyramidal neurons, despite receiving selective LGN inputs at the same strength as other layers, exhibited a much lower OSI due to constant FB currents, which elevated the baseline of their stimulus-evoked responses. Because inhibitory interneurons were not directly modulated by selective external inputs and excitatory-to-inhibitory synapses were much denser than other connections (Suppl. Fig. S1), inhibitory OSI values remained lower in this model configuration, consistent with the results shown in Fig. 2D. In all layers, visualization of single-neuron responses revealed both bimodally tuned pyramidal neurons and more broadly tuned inhibitory neurons (Fig. 3E). The simulation results of realistic OSI and evoked firing rate distribution were maintained for more realistic scenarios. For instance, if we considered that our cortical column model was spatially embedded in a column of radius ∼125 um and translated those quantities to better approximate the values of uniform connection probabilities, as it has been done in our previous work^46^, the distribution remained the same (Suppl. Fig. S3).

### Structural plasticity provides an alternative mechanism for like-to-like connectivity

Our previous results demonstrated that under fine-tuned distributed FF and FB inputs, the cortical column model established by previous studies^40,46^ could replicate the distribution of both OSI and evoked firing rates across laminar layers and cell types. How these OS properties are efficiently transmitted across cortical circuits within the column is an equally important issue. Experimentally, realistic axonal projection pathways originating from the LGN have been reported to follow the so-called canonical microcircuit sequence LGN→ L4→ L2/3→L5→ L6^15,47^. Likewise, the orientation selectivity following this pathway is first observed in L4 and subsequently broadcasted across the column through inter-laminar transmission^15^. This pathway for distributed OS appearance in V1 does not fully align yet with the computational model proposed here so far, which purely relied on highly selective Poisson inputs to all layers (although lateral posterior thalamic nucleus has been found to weakly project to other laminae^45^). Unlike other mammals such as monkeys and cats, which display orientational selectivity maps in their primary visual cortex^2,5,48^, rodent V1 lacks such a map, with differently tuned neurons intermixed in anatomical space. This implies that the PO in rodent V1 is independent of cortical distance. So how could OS in layer 4 be effectively transmitted to other layers without degradation? Recent studies in synaptic physiology suggest that there are co-tuned subnetworks of interconnected cells in the rodent visual cortex^17,18,20,21,49^, indicating that neurons responsive to the same visual feature are more likely to inter-connect, regardless of their spatial distance^20^. These intermingled subnetworks, observed in both vivo and vitro, may correspond to the intermingled ensembles of cells tuned to different orientations.

The synaptic connectome in our baseline cortical column model (model #1) was based on three main neurotransmitter receptors: NMDA, AMPA and GABA_A_ receptors, and the probability of forming a synapse was solely dependent on anatomically recorded average connectivity probability without any connectivity preference in-between pyramidal cells^40^. In order to transform the random baseline connectome into a functional, co-tuned connectome, we implemented a Hebbian type of structural plasticity, which depended on pre- and post-synaptic activities^50^ in a distributed configuration with oriented inputs (Fig. 4A). If pre- and post-synaptic cells are coactive, the synapse was potentiated with probability p+, while if one cell was active and the other inactive, then the synapse was depressed with probability p−. If neither cell was active, no change occurred (Fig. 4A left). Consequently, in the modified model (denoted here as model #2), neurons responsive to the same orientation were more likely to form synapses, while neurons responsive to different orientations tended to lose their established synapses (Fig. 4A right).

**Figure 4:**
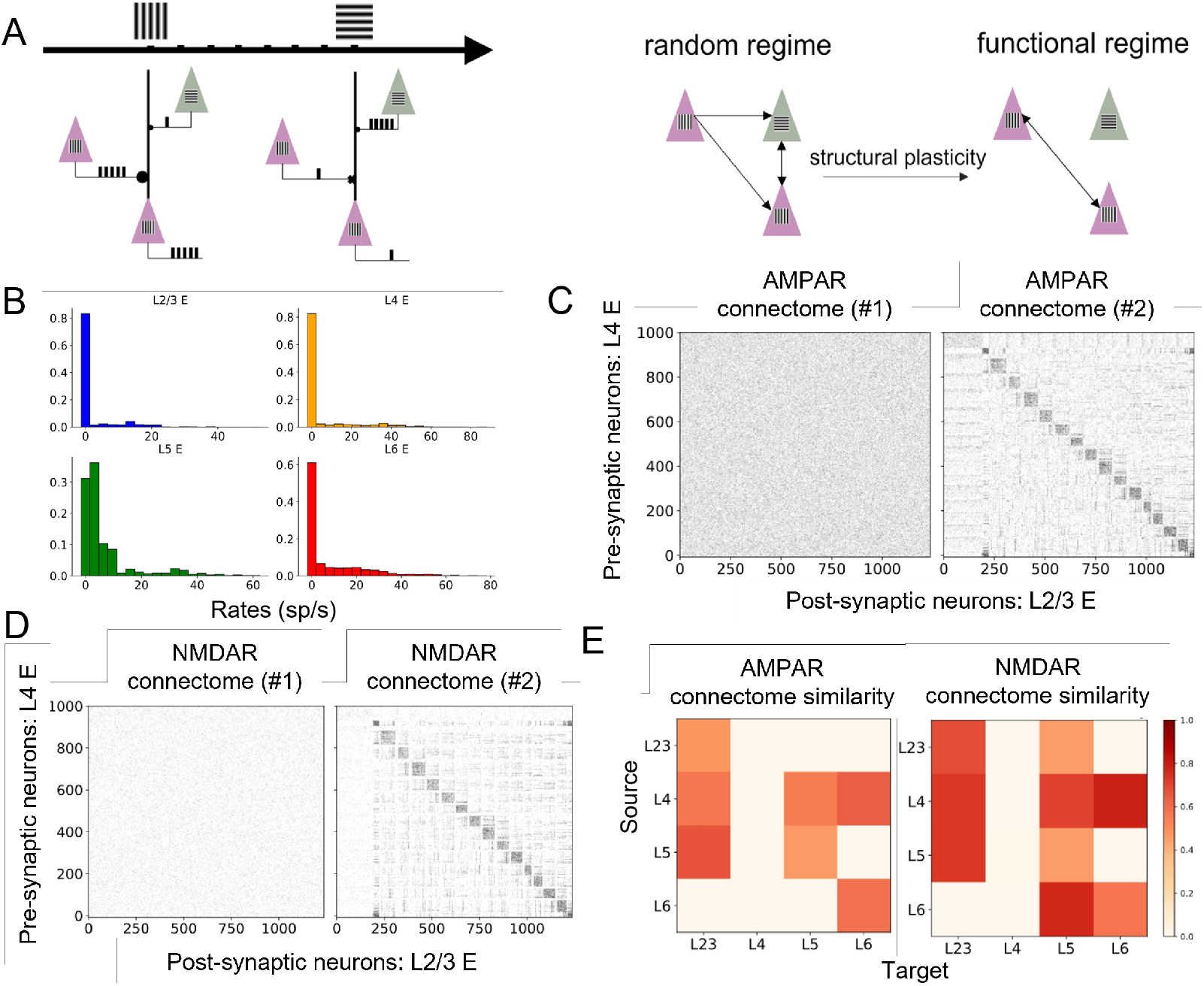
(A) Schematic of the structural plasticity rule. Left: In the distributed configuration, pyramidal neurons respond selectively to visual orientations based on their connectome with the thalamus. For neurons receiving thalamic inputs with the same preferred orientation, the synapse is potentiated (symbolized by <u>a</u> dot) during the preferred orientation period, while remaining unchanged during other orientation periods. For neurons inheriting different preferred orientations, the synapse is depressed (symbolized by <u>a</u> cross) during their individual preferred orientation periods, while remaining unchanged during other orientation periods. Right: After an 8000 ms selective stimulus duration period with 16 different orientations, the initial random connectome transforms into a functional, co-tuned connectome. In this co-tuned connectome, pyramidal neurons with the same orientation preference are more likely to form synapses, whereas pyramidal neurons with different orientation preferences are more likely to delete them. (B) The firing rate distribution of pyramidal cells in the distributed configuration under visual orientation = 0°. (C) Binary L4E- L2/3E AMPAR connectome, black indicates a synapse. Neuron identities have been reordered according to their maximal average firing rates across 16 visual orientations. (D) The same notations as in (C) but for NMDAR connectome. (E) The cosine similarity matrix of the baseline connectome and the modified connectome for AMPA and NMDA receptors.

To determine the threshold for defining a neuron as active, we visualized the pyramidal firing rate distribution (Fig. 4B) in the distributed configuration under a single visual orientation. For excitatory neurons within layer 2/3, 4 and 6, which received only selective Poisson external inputs, the resulting rate distribution displayed an exponential-like distribution. In contrast, for layer 5, due to the modulatory constant FB input, the majority of firing rates clustered around 5 spikes/s (Fig. 4B). Consequently, we set the activity thresholds at 15, 20, 20 and 30 spikes/s for pyramidal cells in layers 2/3, 4, 5, and 6, respectively. We then implemented the structural plasticity described earlier with fine-tuned plasticity parameters for subsequent studies within this paper (see Methods). The biological foundation of this approach is that before the critical period of new-born rodents, visual cortical neurons were intermixed regardless of their laminar locations, so in this sense LGN selectively targets pyramidal neurons in a distributed manner. In the meantime, structural plasticity would be taking place rapidly in the cortex, continuously forming early-stage synaptic connections until the end of the critical period. Next, the neocortex would assume a more definitive shape and, from now on, synaptic connections would be more hard-wired^51–53^.

To better visualize connectome modifications due to plasticity, we reordered neuronal identities according to their maximum average firing rates across visual orientations, and observed that both NMDAR and AMPAR connectomes were significantly modified according to their selective responses, as illustrated in Fig. 4A. Comparing the modified connectome with the original one, synapses between pre- and post-synaptic neurons responsive to the same directions exhibited an increased number of direction-selective synapses, while synapses between neurons responsive to different visual directions were mostly eliminated (Fig. 4C, D). We used cosine similarity as a metric to evaluate the level of structural plasticity in terms of intra- and inter-laminar synaptic turnover, and found that postsynaptic pyramidal cells, in all layers except layer 4, underwent substantial changes (Fig. 4E).

### Functional co-tuned network significantly improves inter-laminar selective transmission

We then explored the laminar responses to direction-selective LGN inputs in the modified model (model #2) presented above. Unlike the distributed configuration in Fig. 3A, this time we modeled LGN Poisson inputs targeting only on layer 4 pyramidal neurons, as suggested by anatomical evidence, and constant FB currents were only injected into layer 5 pyramidal neurons to represent top-down modulation (Fig. 5A).

**Figure 5:**
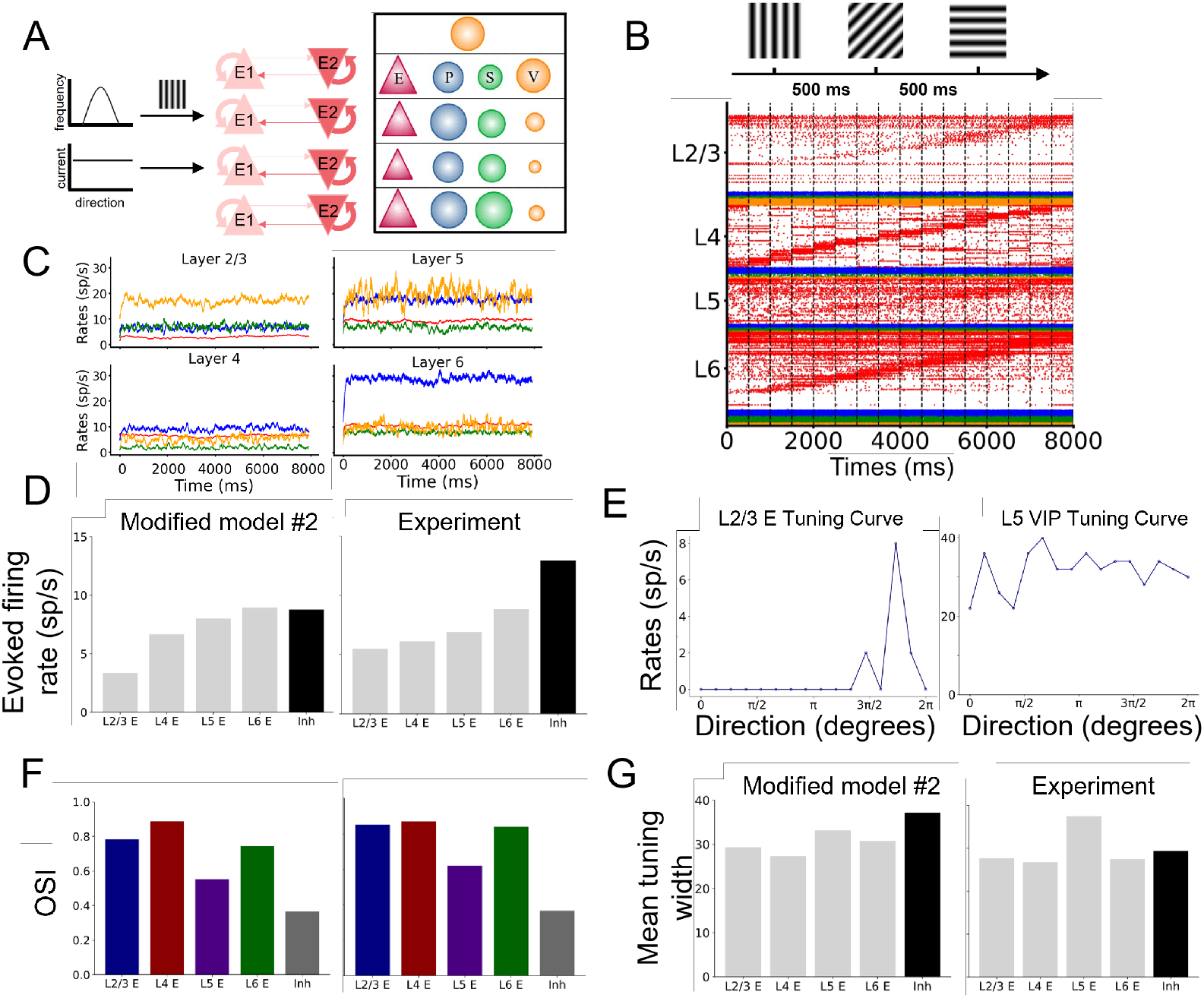
(A) LGN direction-selective inputs project solely to layer 4 pyramidal neurons and FB inputs are injected only into layer 5 pyramidal neurons. (B) Each visual oriented stimulus lasts for 500 ms and firing rates of LGN Poisson neurons follow the Von Mises distribution. Raster plots of the column responses to oriented stimuli, reordered according to individual firing rates at each orientation bin. (C) Mean firing rates of individual cell types during the stimulus duration. (D) Comparison of mean evoked firing rate in the model with experimental values^**1**^. (E) After presenting selective inputs across 16 directions, tuning curves of a L2/3 pyramidal neuron (left) and a L5 VIP neuron (right) were obtained. (F) Comparison of mean OSI in the model with experimental values^**1**^. (G) Comparison of mean tuning width of model subpopulations with experimental values^**1**^.

Following the same simulation protocol as in Fig. 3B, we exposed the modified cortical column model to 16 different visual directions, lasting a total of 8000 ms (Fig. 5B). The resulting spiking activities were similar to the baseline model (model #1), in which pyramidal neurons displayed selective responses to different visual orientations, while interneurons responded broadly without preference (Fig. 5B, C). Furthermore, the average evoked firing rate matched the experimental data^1^ (Fig. 5D), and remained stable across all visual orientations (Fig. 5C). When computing the OSI in the functional co-tuned regime with only layer 4 FF selective input, we found that the model successfully replicated the distribution of OSIs across all layers (Fig. 5F). Two examples, one of a sharply tuned pyramidal neuron and another of a broadly tuned inhibitory neuron are shown in Fig. 5E. In the modified model, cortical OS originating from LGN FF inputs targeted layer 4 pyramidal neurons, and then the information was effectively transmitted to other layers through co-tuned subnetworks, leading to the tuning behavior across the column. Because layer 5 pyramidal cells additionally received constant FB modulation inputs, their baseline activities were significantly higher than excitatory neurons located in other layers, resulting in a low OSI value (Fig. 5F). Besides, the tuning width distribution, defined as the halfwidth at the half-maximal response above baseline^11^ (see Methods), was also consistent with experimental data^1^ except for interneurons (Fig. 5G). In our simulation results, the average tuning width of inhibitory neurons was slightly higher than observed in experiments.

Compared to the distributed configuration in the baseline cortical column model, which included equally strong selective LGN FF inputs to pyramidal cells across all layers and constant FB modulatory inputs to interneurons in deep layers, our modified column model achieved the same effects using only one source of selective LGN inputs and one source of constant FB inputs.

### Perturbation analysis and mean-field analysis uncover the mechanism of evoked firing rates

We further investigated the mechanisms by which like-to-like excitatory intra- and inter-laminar connectomes triggered highly-evoked firing rates (Fig. 5D) compared with the baseline random connectome (model #1; Fig. 2E) under consistent FF and FB configurations. We first noticed that the structural plasticity rule slightly increased the average connection probability between some excitatory laminar populations, especially for the NMDAR connectome (Suppl. Fig. S4C). To analyze the impact of this increase, we took the connection probabilities generated in the modified model and used them to constrain the basal model while keeping the synaptic strength constant (we termed this configuration the random model; or model #3). After simulating the random model using the same protocol as before (Fig. 5A), we calculated the evoked firing rates and found that only L4 E and L5 E were activated, while L2/3 E and L6 E remained silent (Fig. 6A right), indicating that average connectivity probability alone had little effect on laminar excitatory signal propagation in the random model.

**Figure 6:**
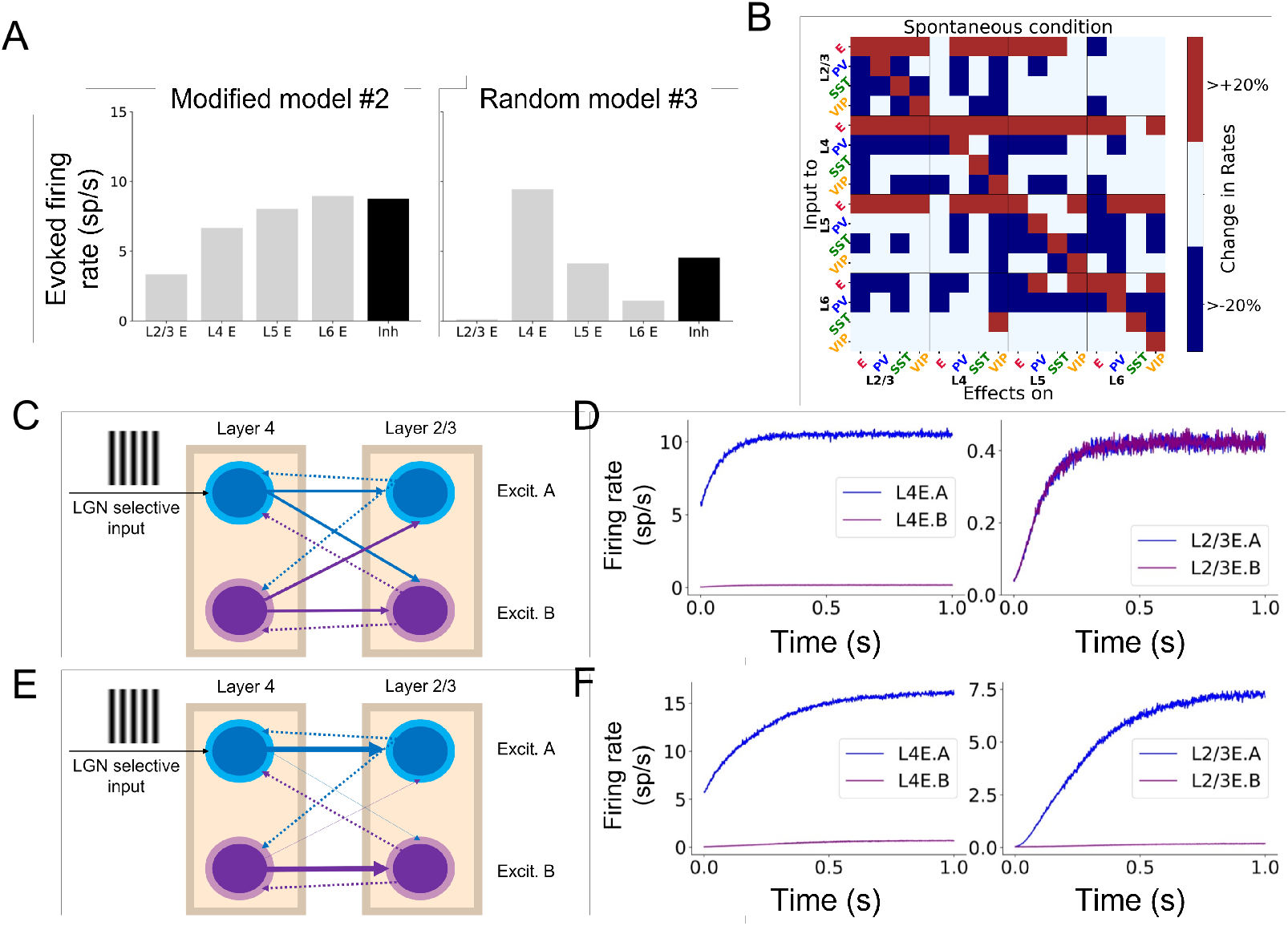
(A) Comparison of mean stimulus-evoked firing rate in the modified model (model #2) and the random model (model #3) with the same FF and FB configuration. (B) Matrix of input-output relationships of the network in the spontaneous state. A detailed description can be found in previous work^**46**^. (C) Mean-field simplification of the layer 2/3-layer 4 microcircuit within our cortical column network (model #3). In each laminar module, cortical dynamics follow a local winner-take-all model, and modules are equally coupled to the same source and target cell types because of the random connection nature of the model^**40**^. Inhibitory neurons are omitted in the schematic but included in the dynamics following a Wong-Wang formalism, and the population L4 E.A is assumed to receive selective LGN inputs under the current orientation. (D) Mean firing rate responses of four populations. Left, layer 4 displays bifurcation because of the external input. Right, differently tuned excitatory populations in L2/3 respond weakly and equally despite fluctuations. (E) Schematic of the mean-field model with the modified connectome (model #2). With similar connection probabilities as in the random connectome, coupling between co-tuned modules increases while coupling between differently tuned modules decrease, leaving their summation to be constant. (F) Mean firing rate responses of four populations. Left, layer 4 displays bifurcation because of external input to L4E.A. Right, layer 2/3 also displays bifurcation because L2/3 E.A now receives stronger FF excitation from L4 E.A than L2/3 E.B does.

Interestingly, compared to the modified model (model #2), the random model showed increased layer 4 excitatory responses to constant orientation stimuli (Fig. 6A). Considering the complexity of recurrent behavior in the laminar cortical column, we adopted here a previously developed perturbation analysis method to explore the modulatory relationship between cell types in response to optogenetics stimulation^46^ (see Methods). To quantify the effects of input perturbations in our network, we defined the response matrix as an array R, in which each component *R*_*XY*_ describes the activity change of population X as a result of slightly increasing the input to population *Y*. We first kept the network in the spontaneous state and then we injected input to one cell group at a time. Next, we observed the effects on the other neuron groups to build the matrix (Fig. 6B). For example, when we perturbed layer 2/3 pyramidal neurons in the spontaneous state, we observed a substantial increase in layer 5 PV activity. When delving into the exact values of the perturbation effects (Suppl. Fig. S6), we found that activating L5 E and L6 E cells caused an overall disynaptic inhibition of L4 E neurons. Consequently, the excitatory transmission originating from layer 4 laterally inhibited its average firing rate responses through disynaptic inhibition from other laminar pyramidal neurons, leading to less activation in the like-to-like connectome scenario.

Finally, we adapted an existing distributed mean-field model^23,54^ to provide further insights into the advantages of like-to-like connectivity on inter-laminar signal propagation. For spiking neurons with ligand-gated ion channels, mean-field analysis can reduce high-dimensional spiking networks into simplified population-level systems based on average firing rates^23^, leading to orientation-selective excitatory populations and non-selective inhibitory populations. We first considered a local circuit consisting of two stimulus-selective excitatory populations and one nonselective inhibitory population as a template to describe simplified neural dynamics within a given layer. For simplicity we focused on the excitatory circuits between layers 4 and 2/3, and since the structural plasticity targeted only E-E connections, we omitted inhibitory populations in the schematic for clarity (Fig. 6C,E). When simulating the random mean-field model with an external current injected into the excitatory layer 4 population selective for stimulus A (termed L4 E.A), we observed a bifurcation in L4 E, as L4 E.B (i.e. the excitatory L4 population selective for stimulus B) received no external input. However, due to the constant intra- and inter-laminar coupling, firing rate responses in L2/3 E were equally low except for some fluctuations caused by background noise (Fig. 6D, right). In contrast, with the like-to-like connectome, excitatory layer 4 populations had a stronger coupling towards the co-tuned putative layer 2/3 populations. As the average number of synapses was kept constant in both scenarios, the total coupling strength between two specific cell types was similar with the random model (e.g. the sum of coupling strengths between L4 E and L2/3 E), so the increase in co-tuned coupling strength (e.g. synaptic strength between L4 E.A and L2/3 E.A, and the one between L4 E.B and L2/3 E.B) led to a decrease in coupling between differently tuned excitatory populations (e.g. L4 E.A-L2/3 E.B, L4 E.B-L2/3 E.A). When stimulating the modified mean-field model with an external current injected into the excitatory subpopulation A in layer 4, bifurcation appeared in both layers 2/3 and 4 due to excitatory signal propagation. Besides this, mean excitatory activity in layer 4 also increased from lateral excitation effects of evoking layer 2/3 excitatory cells (Fig. 6D,F). Furthermore, we can also increase the excitatory self-coupling strength in layer 2/3 to reproduce the facilitation modulation of the intra-laminar like-to-like connectome (Suppl. Fig. S5B), or include strong inter-laminar inhibitory-excitatory coupling to reproduce the disynaptic inhibition of activating deep excitatory populations on excitatory layer 4 activity. Therefore, the simplified mean-field laminar model could quantitively reproduce all phenomena observed in the large-scale spiking network.

## Discussion

We employed columnar-scale spiking network modelling to study the emergence of layer-specific orientation selectivity in mouse V1. To this end, we initially adopted a cortical column model with no connectivity preference across cell types^40^. Under this configuration, direction-selective inputs targeting solely layer 4 pyramidal neurons were not effectively transmitted to other layers. This deviates from state-of-the-art experimental data^1^ reporting that the firing rates evoked in response to drifting grating stimuli were on average above 5 Hz for individual layers. Therefore, the baseline description (or model #1), which followed the known anatomical pathway (LGN → Layer 4 → other layers), failed to reproduce the experimental results. To address this discrepancy, we injected distributed feedforward direction-selective inputs and untuned feedback currents from higher areas to the cortical column. This approach successfully reproduced the experimental OSI and evoked firing rate distribution. However, it also relied on the assumption of random connectivity between pyramidal cells in layers 2/3-6. Recent semi-chronic two-photon calcium imaging recordings, on the other hand, suggest that the axonal projections between pyramidal cell populations in layers 2/3-6 are not random. Instead, they partly rely on the tuning properties of post- and presynaptic neurons, following the like-to-like connectivity of co-tuned subnetworks^20,22^. Motivated by these findings, we implemented a Hebbian structural plasticity rule on pyramidal AMPAR and NMDAR synapses to create a like-to-like connectome. This model with embedded structural plasticity (model #2) was finally able to reproduce the distribution of OSI values and evoked firing rates of pyramidal cells using LGN input solely targeting L4 E and feedback input targeting L5 E, adhering to realistic anatomical pathways.

To gain deeper insights into the advantages of a like-to-like organization, we analyzed a simplified mean-field circuit model with population-coupling strength representing the connectivity preference of two subpopulations. In the scenario of random connections, coupling strength was the same for differently tuned, but same cell-type postsynaptic populations, leading to weak signal propagation and diminishing bifurcation. It should be noted that in this case, the OSI value could be very high as a single neuron could stay silent for most visual orientations but respond weakly to one, resulting in an OSI close to one. In contrast, in the like-to-like connectome, the overall population-coupling strength was attributed to co-tuned coupling, which significantly increased the evoked firing rates in the post-synaptic populations due to structural plasticity. Meanwhile, the winner-take-all behavior induced by non-selective inhibitory neurons led to a strong bifurcation ramping, resulting in high OSI values. This demonstrates that the like-to-like pyramidal-to-pyramidal cell connectivity enhances excitatory transmission, which is crucial for efficient propagation of visual information and orientation selectivity in V1.

Various computational frameworks have been proposed to explain the emergence of orientation selectivity in rodent V1. The traditional view is based on the idea that LGN to V1 axons are untuned, leading to the generation of orientation selectivity in layer 4 pyramidal neurons from the specific spatial projections of LGN neurons^12,14,55,56^. But recent findings support an opposing view, suggesting that cortical orientation selectivity is inherited from LGN-tuned feedforward axons and then amplified within the cortex through recurrent connections^15,16,44^. Consequently, the latest modelling studies have incorporated tuned LGN inputs to explore the corresponding layer-specific behavior^13,57^.

Compared to these recent models, our approach stands out in several ways. For instance, Billeh et al.^13^ used anatomical connectivity data as starting point, but their model later loses this information in favor of an optimization protocol to fit electrophysiological responses. Merkt et al.^57^, on the other hand, nicely captured laminar OSI profiles but failed to capture the evoked firing rate distribution of all cells. As indicated in our Fig. 2, the cortical column model without connectivity preference could only propagate orientation information weakly, leaving average firing rates almost constant. In contrast, our model uses synaptic strengths and connectivity probabilities directly adopted from experimental databases, allowing it to successfully reproduce both OSI and evoked firing rate distributions simultaneously.

Our study still presents some limitations, such as the failure to reproduce significant ramping activities of inhibitory cells (Fig. 5D) and medium tuning width compared to excitatory neurons in layer 4 and other layers^1^. The latest computational models propose the existence of co-tuned subnetworks between excitatory and inhibitory populations, which may be formed during mammalian development^21,58^. Future work could focus on inducing like-to-like connectivity between E-I synapses in a temporally continuous manner and examining whether ramps and medium tuning of I cells can be simulated. This approach would surely increase excitatory transmission to interneurons while decreasing their tuning widths, thus providing a more comprehensive understanding of the mechanisms underlying orientation selectivity in primary visual cortex.

## Materials and methods

The detailed architecture of the model and all parameter specifications are comprehensively described in the methods section of our previous work^40^. Below, we summarize the main features of the model.

### Structure of the baseline model #1

The cortical column model, illustrated in Fig. 1A, comprises 5,000 neurons across five layers. Layers 2/3, 4, 5, and 6 contain pyramidal neurons, PV, SST, and VIP cells, while layer 1 contains only VIP cells. In the four ‘complete’ layers, pyramidal neurons make up 85% of the cells, and inhibitory interneurons 15%. Neuron type proportions in each layer, derived from the Allen Database, are detailed in Supplementary Tables 1 and 2 in Moreni et al.^40^. All neurons receive background noise from the brain, with levels specified in Supplementary Table 3 in Moreni et al.^40^.

Connections refer to pre- and postsynaptic neuron types in each layer, with connection probability (p) indicating the likelihood of a connection. For instance, *p* = 0.1 means a 10% chance of connection between neuron pairs. The connectivity probability matrix *P* includes 288 probabilities across 17 cell groups and is available on the Allen database portal (https://portal.brain-map.org/explore/models/mv1-all-layers). Connection strength varies by neuron group, specified by the matrix *S*, also available online. For example, VIP cells in layer 1 (group X) connecting to SST cells in layer 4 (group Y) have uniform connection strength, though only connect with probability *p*.

### Model for neurons

All pyramidal cells and all three types of interneurons are modelled as leaky integrate-and-fire (LIF) neurons. Each type of cell is characterized by its own set of parameters: a resting potential *v*_*rest*_, a firing threshold *V*_*th*_, a membrane capacitance *C*_*m*_, a membrane leak conductance *g*_*L*_ and a refractory period *τ*_*ref*_. The corresponding membrane time constant is *τ*_*m*_ = *C*_*m*_*/g*_*L*_. The membrane potential *V*(*t*) of a cell is given by:

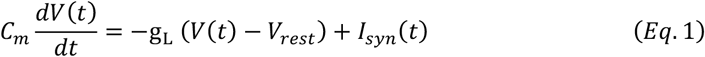

where *I*_*syn*_(*t*) represents the total synaptic current flowing in the cell.

At each time point of simulation, a neuron integrates the total incoming current *I*_*syn*_ (*t*) to update its membrane potential *V*(*t*). When the threshold *V*_*th*_ is reached a spike is generated, followed by an instantaneous reset of the membrane potential to the resting membrane potential *V*_*rest*_. Then, for a refractory period *τ*_*ref*_, the membrane potential stays at its resting value *V*_*rest*_ and no spikes can be generated. After *τ*_*ref*_ has passed, the membrane potential can be updated again (see Tables S4-S8 in Moreni et al.^40^ for the corresponding parameter values).

### Model for synapses

Each cell group in each layer connects to all other groups in the cortical column with its own synaptic strength and probability. Matrix *P* values represent the probability that a neuron in group A (e.g., a PV cell in layer 4) connects to a neuron in group B (e.g., an SST cell in layer 5). Excitatory postsynaptic currents (EPSCs) are mediated by AMPA and NMDA receptors, while inhibitory postsynaptic currents (IPSCs) are mediated by GABA_A_ receptors.

The inputs to model neurons consist of three main components: background noise, external (e.g. sensory) input and recurrent input from within the column. EPSCs due to background noise are mediated in the model exclusively by AMPA receptors *I*_*ext,AMPA*_ (*t*) and EPSCs due to external stimuli (i.e. originating from outside the column) are represented by *I*_*ext*_ (*t*). The recurrent input from within the column is given by the sum of *I*_*AMPA*_ (*t*), *I*_*NMDA*_(*t*), *I*_*GABA*_ (*t*). These are all the inputs from all the other presynaptic neurons projecting to the neuron under consideration.

The total synaptic current that each neuron receives is given by:

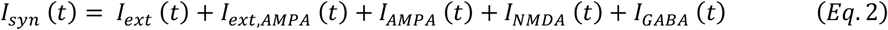

with the last four terms given by:

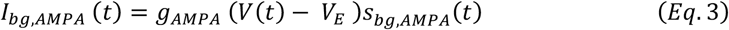

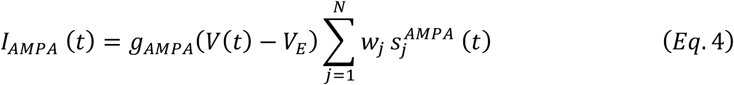

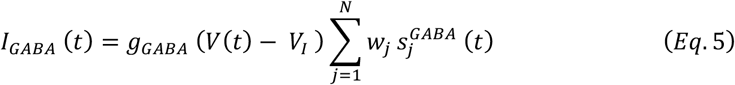

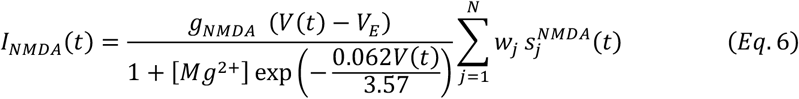

where the reversal potentials are *V*_*E*_ = 0 *mV, V*_*I*_ = *V*_*rest*_, and each group of interneurons has its own *V*_*rest*_. The *g* terms represent the conductances of the specific receptor types. The weights *w*_*j*_ represent the strength of each synapse received by the neuron. The sum runs over all presynaptic neurons *j* projecting to the neuron under consideration. NMDAR currents have a voltage dependence controlled by extracellular magnesium concentration, [*Mg*^2+^]=1 mM. The *s* terms represent the gating variables, or fraction of open channels and their behaviour is governed by the following equations.

First, the AMPAR channels are described by:

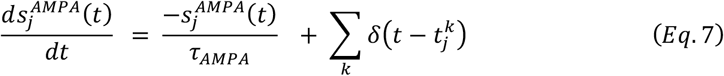

where the decay time constant of the AMPA currents is *τ*_*AMPA*_ = 2 *ms*, and the sum over *k* represents the contribution of all spikes (indicated by delta, *δ*) emitted by presynaptic neuron *j*. In the case of external AMPA currents (Eq. 3), the spikes are emitted accordingly to a Poisson process with rate *υ*_*bg*_. Each group of cells in each layer is receiving a different Poisson rate of background noise (see Table S9 in Moreni et al.^40^). *υ*_*bg*_ is cell-specific, hence each has a specific firing pattern.

The gating of single NMDAR channels is described by:

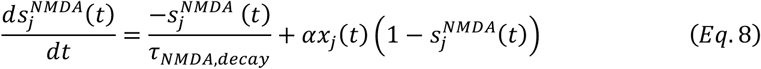

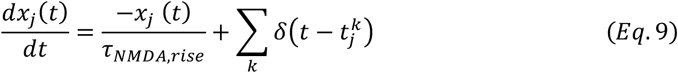

where the decay and rise time constants of NMDAR current are 80 *ms* and 2 *ms* respectively, and the constant *α* = 0.5 ms^−1^. The GABA_A_ receptor synaptic variable is described by:

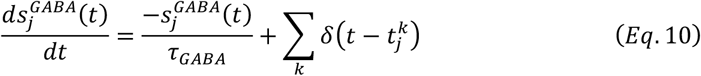

where the decay time constant of GABA_A_ receptor current is 5 *ms*.

### Parameters of the model

As previously mentioned, each type of cell in each layer is characterized by its own set of parameters: a resting potential *V*_*rest*_, a firing threshold *V*_*th*_, a membrane capacitance *C*_*m*_, a membrane leak conductance *g*_*L*_ and a refractory period *τ*_*ref*_.

These data are taken from the Allen institute database (https://portal.brain-map.org/explore/models/mv1-all-layers). For each type of cell in each layer the Allen database proposes different subsets of cells (e.g. two different PV subsets in L4), each with his own set of parameters. To simplify the model, for each layer we only used one set of parameters for each cell type, choosing the set of parameters of the most prevalent subset of cells. The parameters *C*_*m*_, *g*_*L*_, *τ*_*ref*_, *V*_*rest*_, *V*_*th*_ used for each cell type in each layer are reported in Tables S4-S8 in Moreni et al.^40^.

### Details on the background noise

All neurons received background noise, representing the influence of the ‘rest of the brain’ on the target area, as shown in Figure 1. Excitatory postsynaptic currents (EPSCs) due to background noise are exclusively mediated in the model by AMPA receptors, denoted as *I*_*ext,AMPA*_ (*t*) (Eq. 3). The levels of background noise that each group of cells received can be found in Supplementary Table 3 in Moreni et al.^40^. The firing rate *υ*_*bg*_ of the background Poissonian pulse generators connected to each group differed among them.

Each neuron in every group is connected to its own background Poisson generator. Thus, even though the rates *υ*_*bg*_ of the Poisson generators are the same for all neurons within the same group, each specific cell receives its own different pulse train.

### Details on the weights

The weight of each synapse *w*_*j*_ from neurons of group A to neurons in group B is chosen to be equal to

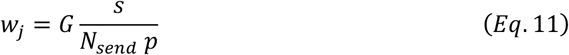

where *G* = 5 is the global coupling factor, *s* is the overall strength between the two connected groups of cells, *N*_*send*_ is the number of neurons in the sending population A, and *p* the probability of connection between the neurons of the two groups (A and B) taken from the experimental probability matrix P. For each pair of connected groups, *s* is taken from the experimental synaptic connectivity matrix S, defined by the 16 ∗ 16 + 16 ∗ 2 = 288 synaptic strengths between the 17 considered cell types (4 groups in each of the 4 layers plus 1 group in layer 1). Matrices *S* and *P* can be found at https://portal.brain-map.org/explore/models/mv1-all-layers. The normalization in Eq. 11 above guarantees that the dynamics and equilibrium points of the system scale properly with the size of the network (Fig. S2).

The spikes generated by an excitatory neuron can target AMPA or/and NMDA receptors of the postsynaptic receiving neuron, with a ratio of 0.8 to 0.2, respectively. Thus, the probability of connection *p* between the neurons of the two groups (A and B) is multiplied by 0.2 for NMDA receptors and 0.8 for AMPA receptors.

For example, suppose excitatory neurons in group A are connected to neurons in group B with a probability *p* (from the matrix P), then the excitatory connections targeting the AMPA receptors of group B will be chosen with *p*_*AMPA*_ = *p* ∗ 0.8 and for NMDA receptors is *p*_*NMDA*_ = *p* ∗ 0.2. All differential equations were numerically solved using Euler’s method, using 0.1 ms time step.

### Selective LGN activities

LGN neurons were modelled as Poisson neurons. For simplicity, we considered a ring topology, so that the visual orientation was parametrized by an angle *ϕ* ∈ [0, 2*π*]. The firing rate of spatially selective neurons was modulated according to a Von Mises distribution:

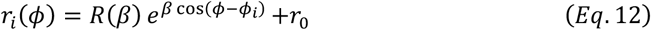

where *ϕ*_*i*_ was the preferred orientation of neuron *i, r*_0_ a baseline firing rate, and 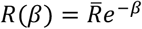 where 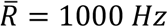 was a constant that set the maximal firing rate. The parameter *β* determined the sharpness of the tuning curve.

### Orientation selectivity index (OSI)

The definition of OSI in our manuscript was consistent with the experimental paper^1^, based on the neuronal firing rates at the preferred and orthogonal orientation:

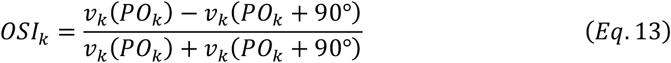

where *v*_*k*_ was the firing rate of neuron *k*, and preferred orientation (PO) was defined arg_θ_ max *v*_*k*_ (*θ*).

### Distributed configuration

In the distributed FF and FB configuration (Fig. 3A), our baseline cortical column model received distributed LGN selective FF inputs targeting L2/3 E, L4 E, L5 E and L6 E, for each layer, the number of LGN Poisson neurons was the same as the corresponding pyramidal population size, and the connectivity probability was *p* = 0.0015, 0.0015, 0.0015 and 0.003 respectively. For FB modulatory constant currents, they targeted on all inhibitory populations in layer 1, 5 and 6, with the value of 30 *pA*.

### Structural plasticity

The plasticity rule was already described in the main text, here we revisited its details: If pre- and post-synaptic cells were coactive, the synapse was potentiated with probability p+, while if one cell was active and the other inactive, then the synapse was depressed with probability p−. The exact parameters in our model were as follows:

**Table 1.** Plasticity values of potentiation for AMPA connectome.

| Source\Target | L2/3 E | L4 E | L5 E | L6 E |
| --- | --- | --- | --- | --- |
| L2/3 E | 0.4 | 0 | 0 | 0 |
| L4 E | 0.4 | 0 | 0.25 | 0.2 |
| L5 E | 0 | 0 | 0.2 | 0 |
| L6 E | 0 | 0 | 0 | 0.2 |

**Table 2.** Plasticity values of depression for AMPA connectome.

| Source\Target | L2/3 E | L4 E | L5 E | L6 E |
| --- | --- | --- | --- | --- |
| L2/3 E | 0.4 | 0 | 0 | 0 |
| L4 E | 0.4 | 0 | 0.4 | 0.4 |
| L5 E | 0 | 0 | 0.4 | 0 |
| L6 E | 0 | 0 | 0 | 0.4 |

**Table 3.**
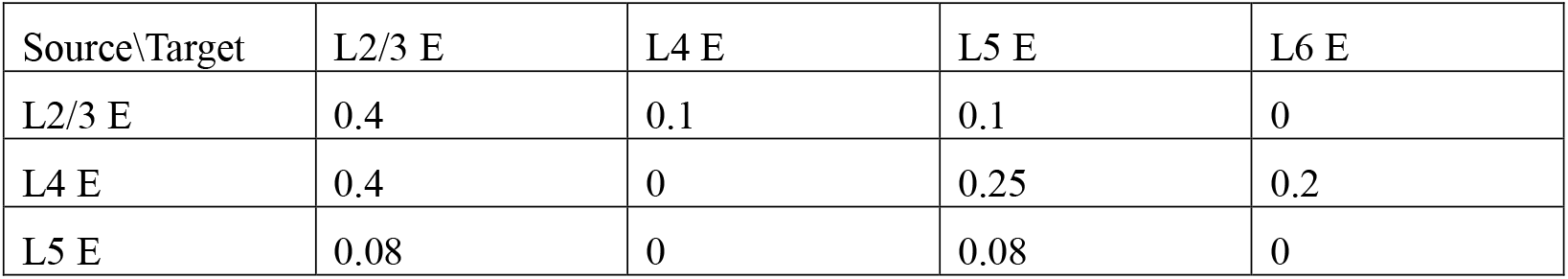
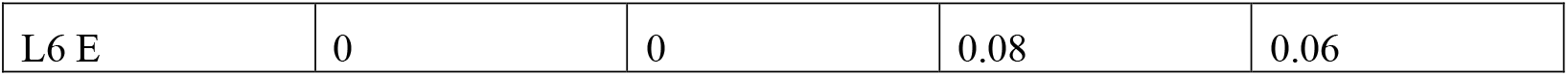
Plasticity values of potentiation for NMDA connectome.

**Table 4.** Plasticity values of depression for NMDA connectome.

| Source\Target | L2/3 E | L4 E | L5 E | L6 E |
| --- | --- | --- | --- | --- |
| L2/3 E | 0.4 | 0.2 | 0.2 | 0 |
| L4 E | 0.4 | 0 | 0.4 | 0.4 |
| L5 E | 0.4 | 0 | 0.3 | 0 |
| L6 E | 0 | 0 | 0.4 | 0.3 |

### Perturbation analysis

We analysed the effects of specific inputs on neuronal populations using matrices (Fig. 6B). Each square in the matrix represents a distinct simulation. The y-axis denotes the population receiving the input, while the x-axis shows the impact on another population. For each simulation, a 30 *pA* constant current was injected into each group, and we recorded the resulting effects on all other neuronal groups. The matrices were constructed based on the percentage change in average firing rate, comparing mean rates after stimulation with baseline rates (prior to input injection). Baseline rates were determined from a 3-second simulation window, which was also used to assess rates after current injection. Red squares indicate a firing rate increase of 20% or more relative to baseline, blue squares indicate a decrease of 20% or more, and white squares indicate changes less than 20%, whether increase or decrease.

### Mean-field model: an example of the L4-L2/3 circuit

We used the model configurations in Fig. 6C-E as an example and firstly employed the Wong-Wang model^23^ to characterize the neural dynamics of a local microcircuit representing V1 L4. This model, in its three-variable version, captures the temporal evolution of the firing rates of two input-selective excitatory populations, as well as the firing rate dynamics of an inhibitory population representing the comprehensive coupling in-between all interneuron types. The populations are interconnected with each other, and the model is governed by the following equations:

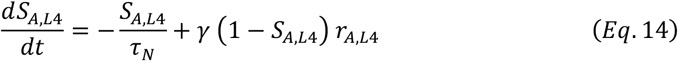

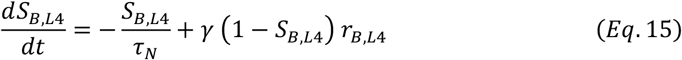

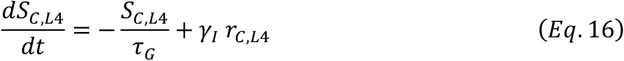

Here, S_A,L4_ and S_B,L4_ are the NMDA conductance of selective layer 4 excitatory populations A and B respectively, and S_C,L4_ is the GABAergic conductance of the inhibitory population. Values for the constants are τ_N_=60 ms, τ_G_=5 ms, γ=1.282 and γ_I_=2. The variables r_A,L4_, r_B,L4_ and r_C,L4_ are the mean firing rates of the two excitatory and one inhibitory population, respectively.

They are obtained by solving, at each time step, the transcendental equation *r*_*i*_ = *ϕ*_*i*_ (*I*_*i*_) (where *ϕ* is the transfer function of the population, detailed below), with I_i_ being the input to population ‘i’, given by

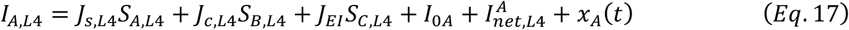

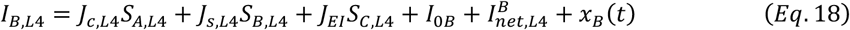

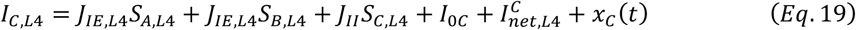

In these expressions, J_S,L4_, J_C,L4_ are the self- and cross-coupling between excitatory populations, respectively, J_EI_ is the coupling from the inhibitory populations to any of the excitatory ones, J_IE_ is the coupling from any of the excitatory populations to the inhibitory one, and J_II_ is the self-coupling strength of the inhibitory population. The parameters I_0i_ with i=A, B, C are background inputs to each population. Fixed parameters across the cortex are J_EI_=-0.31 nA, J_II_=-0.12 nA, I_0A_= I_0B_= 0.25 nA and I_0C_=0.26 nA.

The last term x_i_(t) with i=A, B, C is an Ornstein-Uhlenbeck process, which introduces some level of stochasticity in the system. It is given by

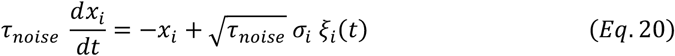

Here, ξ_i_(t) is a Gaussian white noise, the time constant is τ_noise_=2 ms and the noise strength is σ_A,B_=0.01 nA for excitatory populations and σ_C_=0 for the inhibitory one.

The transfer function *ϕ*_*i*_(*t*) which transform the input into firing rates takes the following form for the excitatory populations^40^:

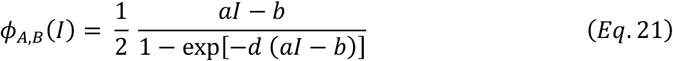

The values for the parameters are *a*=135 Hz/nA, *b*=54 Hz and *d*=0.308 s. For the inhibitory population a similar function can be used, but for convenience we choose a threshold-linear function:

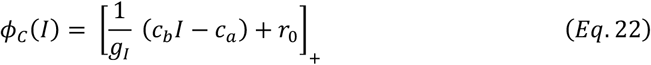

The notation [*x*]_+_ denotes rectification. The values for the parameters are g_I_=4, c_b_=615 Hz/nA, c_a_=177 Hz and r_0_=5.5 Hz.

When the inter-laminar transmission is taken into consideration, inter-laminar currents take the form as:

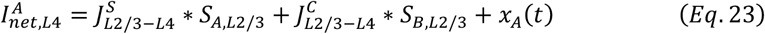

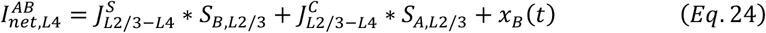

here we take L2/3 E to L4 E coupling as an example, where J_S,L2/3-L4_, J_C,L2/3-L4_ are the self- and cross-coupling between excitatory populations from L2/3 E to L4 E, indicating the inter-laminar coupling. In the case of random model (Fig. 6C), J_S,L2/3-L4_=J_C,L2/3-L4_=0.1nA, J_S,L4-L2/3_=J_C,L4-L2/3_=0.15nA. While in the case of co-tuned model (Fig. 6E), J_S,L4-L2/3_=0.3nA, J_C,L4-L2/3_=0.0nA.

## Acknowledgments

This work was done with the support of EBRAINS and HBP computing services.

## Author contributions

LZ and JFM designed the study; GM provided simulation and analysis code; LZ performed the research; LZ, GM, and JFM analysed and discussed the results; LZ, GM CMAP and JFM wrote the manuscript.

## Data and materials availability

All information needed to reproduce the results of this manuscript are in the main text and Materials and Methods section, and the code used will be made available upon publication of this work.

## Supplementary figures

**Supplementary Figure 1:**
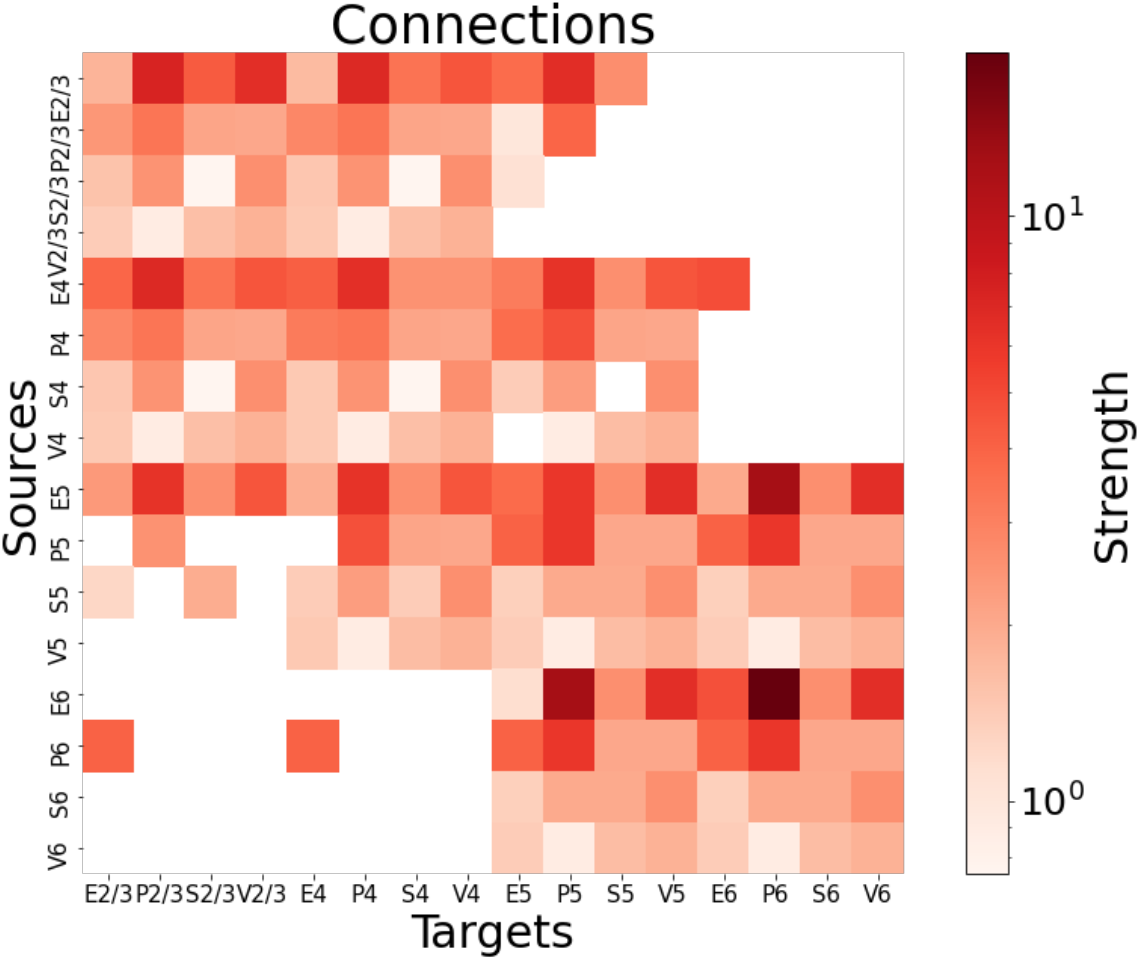
The connectivity matrix of all cell types (adapted from previous work^13^).

**Supplementary Figure 2:**
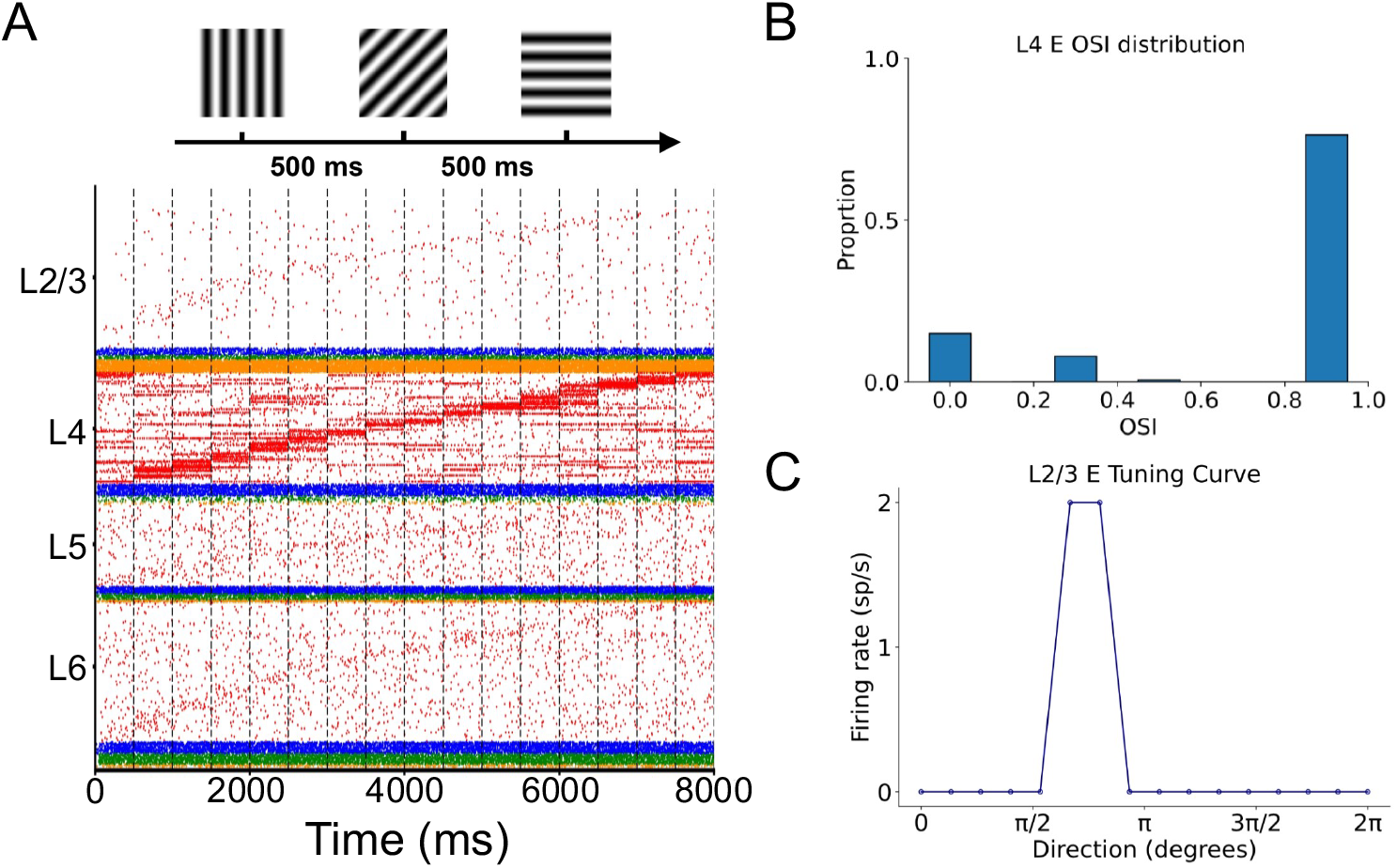
Baseline model with selective L4 input. (A) Each visual orientation lasts for 500 ms and firing rates of LGN Poisson neurons follow the Von Mises distribution. Raster plots of the column responses to orientational stimuli, reordered according to individual firing rates at each orientation bin. (B) The OSI distribution of layer 4 pyramidal neurons. (C) The tuning curve of a layer 2/3 excitatory neuron, with zero responses in all directions except significantly weak responses in the preferred orientation.

**Supplementary Figure 3:**
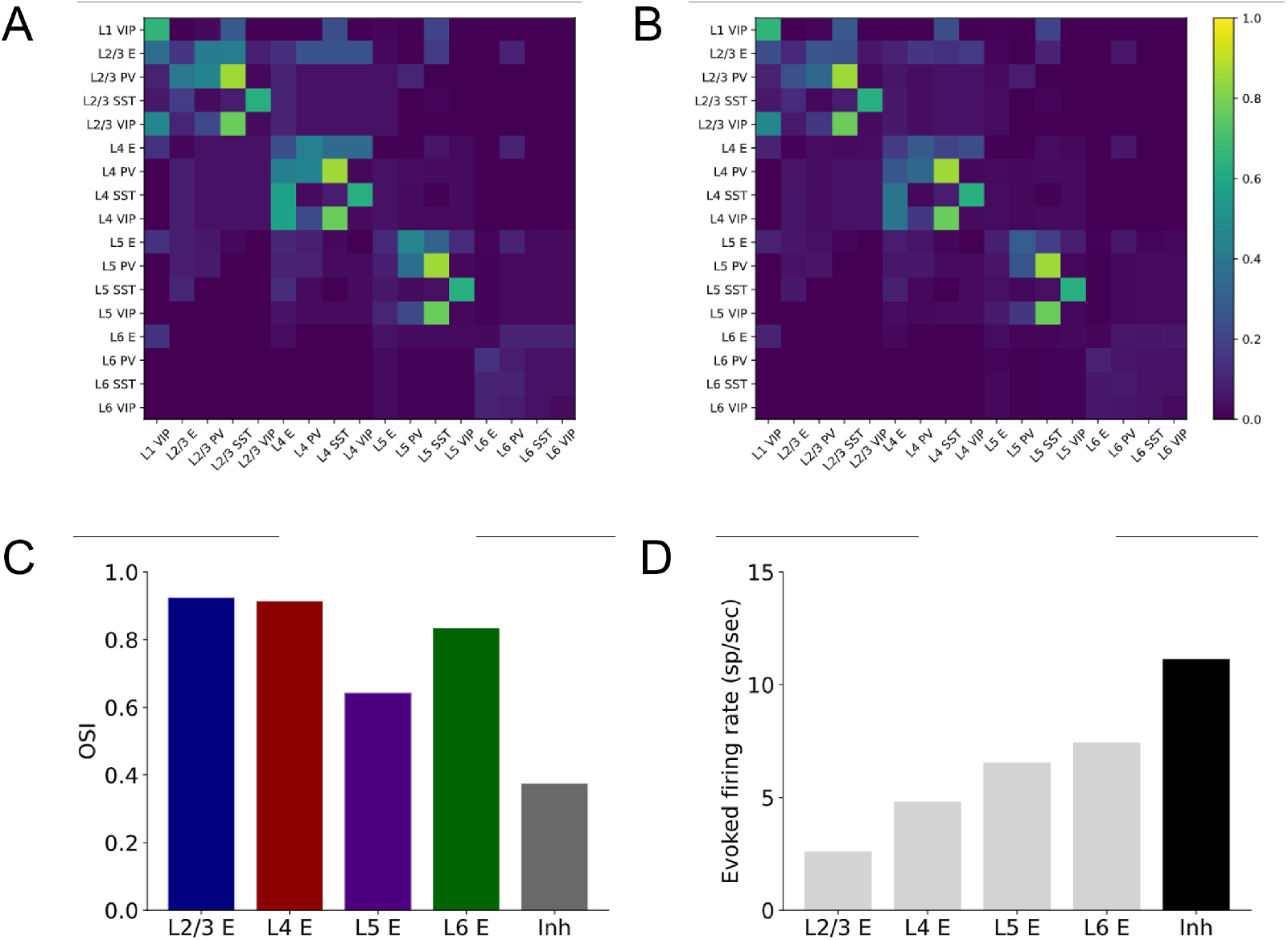
Considering the radius of the column. (A) The original probability matrix. (B) The new probability matrix taking the radius of the column into consideration and translating those quantities to better approximate values of uniform connection probabilities. (C-D) The OSI and evoked firing rate distribution when using the new probability matrix with distributed FF and FB inputs as in Fig. 3.

**Supplementary Figure 4:**
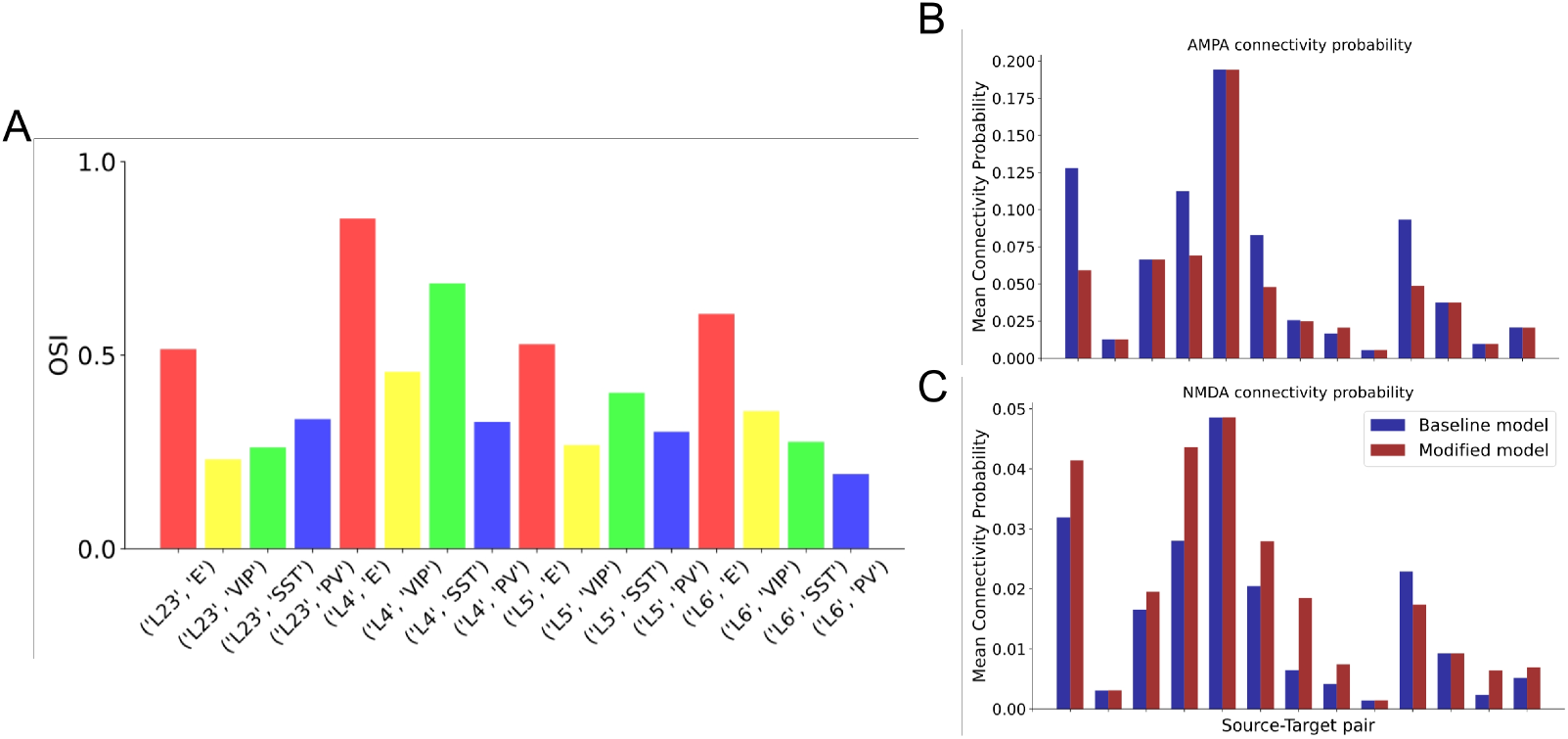
The modified model. (A) OSI distribution with different interneuron cell types. (B)-(C) The comparison of mean connectivity probability between the baseline connectome and the modified connectome, each of them corresponds to one specific source-target excitatory population pair.

**Supplementary Figure 5:**
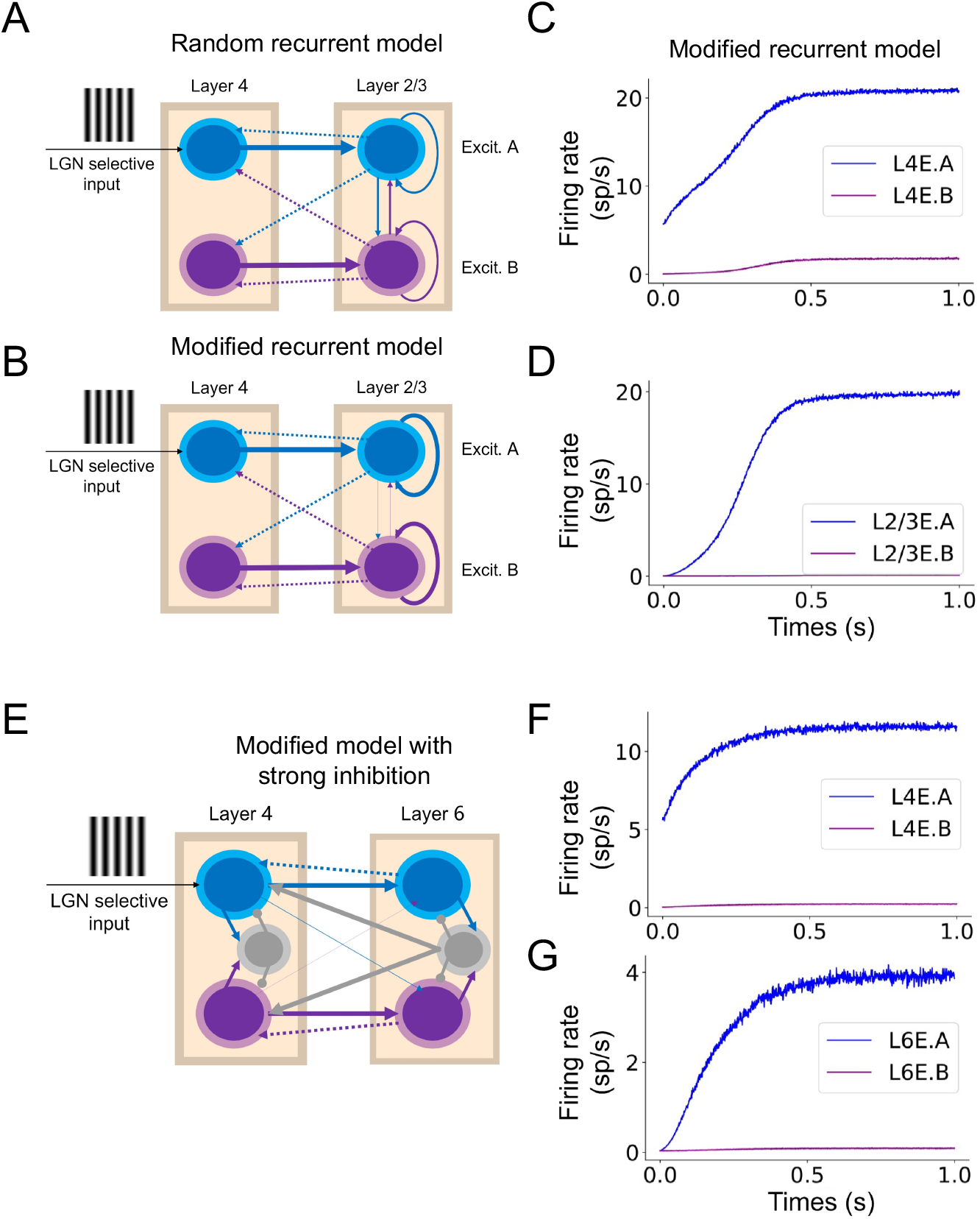
Mean-field analysis on inter-laminar modulation. (A) The same as Fig. 6C, but with the visualization of recurrent excitatory coupling in-between L2/3 E. In the scenario of random projection, the coupling between putatively co-tuned L2/3 E and differently tuned L2/3 E are the same (as indicated by thickness). (B) The same as (A), but in the scenario of the modified model, with higher co-tuned coupling than differently tuned coupling in-between L2/3 excitatory populations. (C)-(D) Mean firing rate responses of four populations. Because of higher self (co-tuned) recurrent coupling, mean responses of L2/3 E.A are higher than the random case in Fig. 6F. (E) The mean-field model can also explain the disynaptic inhibition of L6 (5) E on L4 E. In this case, we considered a strong inhibitory coupling between L6 I and L4 E in the modified configuration. When activating L4 E.A, bifurcation takes place in both L4 E and L6 E because of strengthened co-tuned L4-L6 E coupling. And the activation of L6 E would activate L6 I and this inhibitory activation would finally provide non-preference inhibition on L4 E. Since the coupling between L6 I and L4 E is strong, this inhibition will decrease firing rate responses in L4 E, mainly on L4 E.A.

**Supplementary Figure 6:**
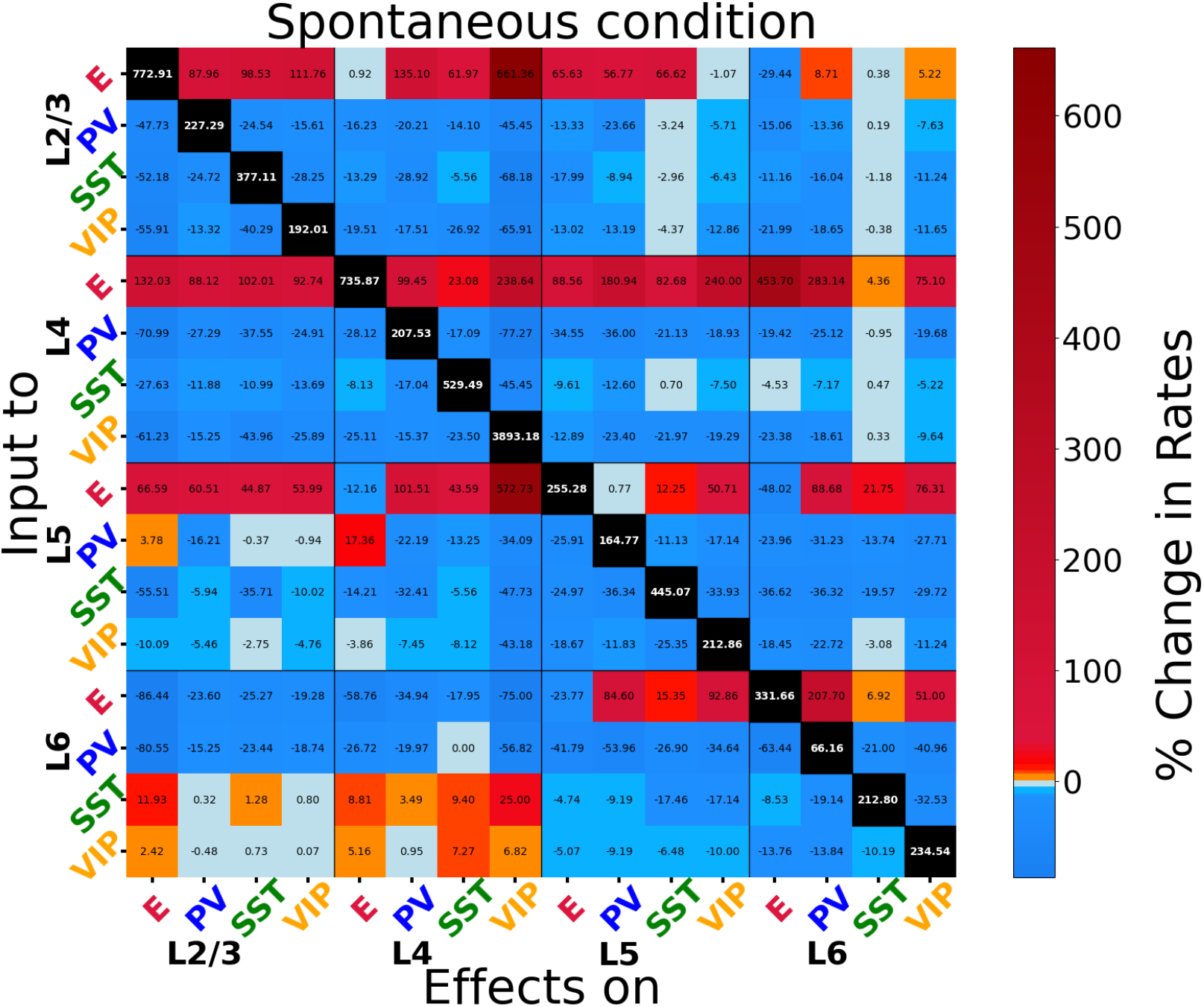
Exact values of the spontaneous perturbation matrix in Fig. 6B.

